# Blood Transcriptional Correlates of BCG-Induced Protection Against Tuberculosis in Rhesus Macaques

**DOI:** 10.1101/2022.11.14.516343

**Authors:** Yiran E. Liu, Patricia A. Darrah, Joseph J. Zeppa, Megha Kamath, Farida Laboune, Daniel C. Douek, Pauline Maiello, Mario Roederer, JoAnne L. Flynn, Robert A. Seder, Purvesh Khatri

## Abstract

Blood-based correlates of vaccine-induced protection against tuberculosis (TB) are urgently needed. We analyzed the blood transcriptome of rhesus macaques immunized with varying doses of intravenous (IV) BCG followed by *Mycobacterium tuberculosis* (*Mtb*) challenge. We used high-dose IV BCG recipients for “discovery” and validated our findings in low-dose recipients and in an independent cohort of macaques receiving BCG via different routes. We identified seven vaccine-induced gene modules, including an innate module (module 1) enriched for type 1 interferon and RIG-I-like receptor signaling pathways. Module 1 on day 2 post-vaccination was highly correlated with lung antigen-responsive CD4 T cells at week 8 and with *Mtb* and granuloma burden following challenge. Parsimonious signatures within module 1 at day 2 post-vaccination predicted protection following challenge with AUROCs ≥ 0.91. Together these results indicate that the early innate transcriptional response to IV BCG in peripheral blood may provide a robust correlate of protection against TB.

## INTRODUCTION

Tuberculosis (TB) persists as a major contributor to global morbidity and mortality, with approximately 10 million new cases and 1.5 million deaths each year [1]. The only available TB vaccine, BCG, is widely administered at birth by the intradermal (ID) route but confers variable durable protection against adolescent and adult pulmonary TB [2; 3; 4]. An understanding of the vaccine-induced immune correlates of protection will guide future TB vaccine development.

Prior studies have focused on adaptive immune responses, especially antigen-specific polyfunctional CD4 T cells [5; 6] which have been shown in many animal models to be critical for vaccine-elicited protective immunity against *Mycobacterium tuberculosis* (*Mtb*). However, in humans vaccinated with BCG or a previously unsuccessful vaccine candidate MVA85A, CD4 T cell responses have failed to correlate with protection [7; 8; 9; 10]. In one study, CD4 T cell responses were associated with TB disease risk but showed insufficient predictive accuracy; in fact, none of the 22 pre-specified immune variables evaluated in this study achieved areas under the receiver operating characteristic (AUROCs) over 0.66 [11]. Thus, it is important to develop models of vaccination that confer high-level protection and use analytical methods that more fully encompass the range of innate and adaptive responses to define more accurate correlates of protection.

Several new models of TB vaccination and challenge have achieved levels of protection which may enable more comprehensive approaches to identifying correlates of protection [12; 13; 14]. Notably, a recent study showed that intravenous delivery of BCG (IV BCG) conferred sterilizing protection against *Mtb* challenge in highly susceptible rhesus macaques [12]. Compared to ID or aerosol (AE) administration of BCG at the same dose, IV BCG elicited higher frequencies of systemic and lung-localized antigen-responsive CD4 and CD8 T cells, as well as increased counts of NK cells, MAIT cells, and Vγ9+ γδ T cells. In addition, IV BCG induced high antigenspecific antibody titers, including IgM, in plasma and bronchoalveolar lavage (BAL) [15]. While these findings suggest potential mechanisms by which IV BCG vaccination mediates protection compared to other routes, the high level of protection among IV BCG-vaccinated animals precluded identification of correlates of protection within this group. Moreover, in this study there was no assessment of blood transcriptional responses to vaccination, which we and others have shown to be important for identifying biomarkers of vaccine immunogenicity or efficacy against other infections including influenza, Ebola, and yellow fever [16; 17; 18; 19]. Such transcriptomic analysis by our group has also provided a blood-based correlate of risk for TB in humans that continues to be independently validated and is actively progressing toward clinical translation [20; 21; 22].

In the present study, we profiled the blood transcriptome in a new cohort of rhesus macaques that were vaccinated with varying doses of IV BCG to achieve a wider range of immune responses and outcomes following *Mtb* challenge. This “dose” cohort is detailed in a separate manuscript (Darrah *et al*, submitted). Here, we applied a systems vaccinology approach to comprehensively assess the blood transcriptional response to IV BCG across several timepoints following vaccination. We hypothesized that the blood transcriptional response would vary by dose, but that distinct blood transcriptional signatures elicited by IV BCG vaccination would nonetheless predict localized immune responses in the lung and correlate with protection following *Mtb* challenge across dose groups. We then sought to validate these signatures in an independent cohort of macaques from the previously published study (“route” cohort) [12] to assess the generalizability of our signatures to other routes of BCG vaccination.

## RESULTS

Thirty-four rhesus macaques of Indian origin were vaccinated with half-log increasing doses of IV BCG (between 3.9×10e4 and 2.5×10e7 CFU) and challenged with *Mtb* twenty-four weeks later (**Fig 1A**). Bulk RNA sequencing was performed on whole blood samples collected at baseline (pre-vaccination) and at two days, two weeks, four weeks, and twelve weeks following IV BCG vaccination (**Fig 1A**). All timepoints were prior to *Mtb* challenge. We were underpowered to perform high-dimensional multivariate analyses adjusting for time, vaccine dose, and protection outcome. However, we hypothesized that vaccine-induced and protection-associated responses would be most evident with high-dose IV BCG and could then be refined with low-dose IV BCG. Therefore, we analyzed high-dose (>10^6^ CFUs of BCG; n=16) and low-dose (<10^6^ CFUs of BCG; n=18) recipients separately as “discovery” and “validation” subsets, respectively (**Methods)**. Finally, we used blood transcriptional data from macaques in the route cohort as further independent validation [12].

**Figure 1.**
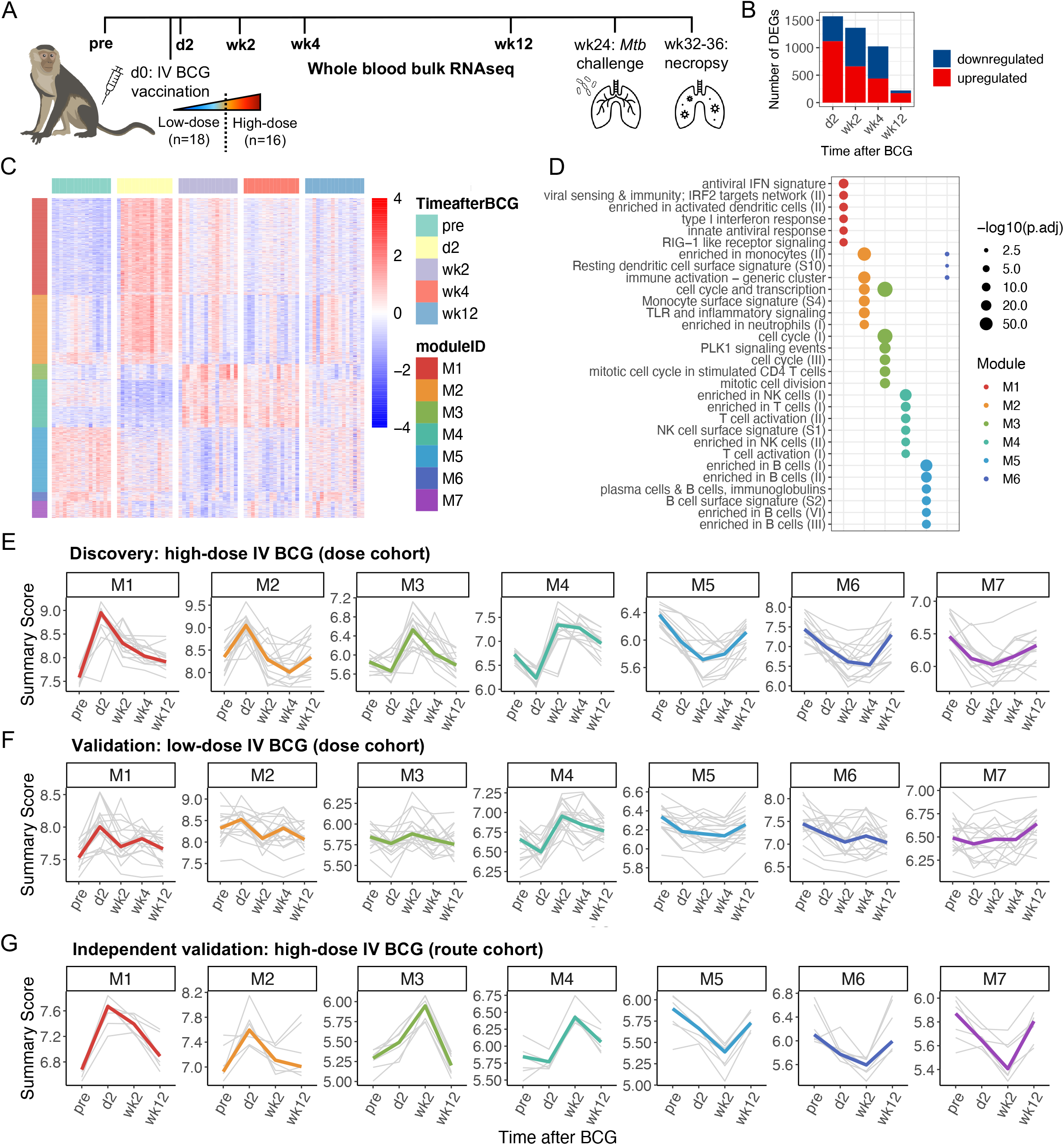
Blood transcriptional response to high-dose IV BCG vaccination. A) Schematic of dose study. Whole blood samples were collected at bolded timepoints for RNA sequencing. Macaques were classified as high-dose IV BCG recipients (N=16) or low-dose IV BCG recipients (N=18). B) Number of upregulated (red) or downregulated (blue) differentially expressed genes (DEGs) in high-dose IV BCG recipients at each timepoint relative to baseline. C) Heatmap of DEGs in high-dose recipients. Expression is scaled per row. Genes are organized by module. Unassigned genes (N=191) are not shown. D) Significantly enriched (adjusted p-value ≤ 0.01) immune pathways within each module. A maximum of six enriched pathways (ranked by significance) are shown for each module. E-G) Summary scores representing the activity of each of the seven modules for individual macaques (thin grey lines) and median responses (thick colored lines) among E) high-dose recipients from the dose cohort, F) low-dose recipients from the dose cohort, and G) high-dose recipients from the independent route cohort. Animals from the route cohort were not sampled at week 4.

In the discovery subset of macaques that received high-dose IV BCG, 2,499 genes were differentially expressed (log2 fold change ≥ 1.5, FDR ≤ 10%) at one or more timepoints relative to baseline (**Fig 1B-C**). Note that we did not consider outcomes following *Mtb* challenge when identifying these differentially expressed genes (DEGs) in order to first analyze the overall response to IV BCG vaccination. Weighted gene correlation network analysis (WGCNA) of these 2,499 DEGs identified seven gene modules (**Fig 1C**). We characterized each module using pathway analysis, with a reference gene set database specifically derived from studying blood transcriptional responses [23]. Module 1 and 2 were upregulated early (day 2 post-vaccination) and enriched for dozens of innate immune pathways including those related to type 1 interferon (IFN) signaling, toll-like receptor (TLR) and RIG-I-like receptor (RLR) signaling, dendritic cell (DC) and monocyte activation, and neutrophil recruitment (**Fig 1D, Table S1)**. Module 3 and 4 were upregulated later (2-4 weeks post-vaccination) and were enriched for pathways involved in CD4 T cell proliferation and T and NK cell activation. Finally, module 5, 6, and 7 were downregulated following vaccination, with module 5 enriched for several B cell pathways and module 6 for a handful of DC and monocyte pathways. No pathways were significantly overrepresented in module 7, even when using a different reference gene set database, suggesting that it may be comprised of genes with unknown roles in the immune response to vaccination.

We summarized the activity of the seven modules over time using a score defined as the geometric mean expression of all genes in each module (**Fig 1E)**. In the validation subset, low-dose IV BCG recipients exhibited similar trends in module activity over time as the discovery subset of high-dose IV BCG recipients (**Fig 1F**). However, the magnitude of vaccination-induced changes in module scores in low-dose recipients was lower and more variable. In extended validation, the activity of all seven modules was recapitulated in high-dose IV-vaccinated macaques from the independent route study (**Fig 1G**). For animals in the previous study that received BCG via other routes, changes in module scores over time were considerably dampened or even undetectable (**Fig S1**).

McCaffrey *et al*. recently demonstrated high concordance between localized immune responses in lung and systemic immune responses to *Mtb* infection in peripheral blood [24]. Further, in studies of vaccines against other bacterial and viral infections, blood transcriptional signatures measured just 1-3 days following vaccination correlated with subsequent antibody responses [18; 23]. Therefore, we investigated whether blood transcriptional responses to IV BCG were correlated with immunological markers in BAL samples collected 4-8 weeks following vaccination (**Fig 2A)**. These BAL features were examined by flow cytometry or Luminex independently of all blood transcriptomic analyses (Darrah et al, submitted). Specifically, we hypothesized that early transcriptional responses to IV BCG in peripheral blood would be associated with subsequent adaptive responses, including those in lung.

**Figure 2.**
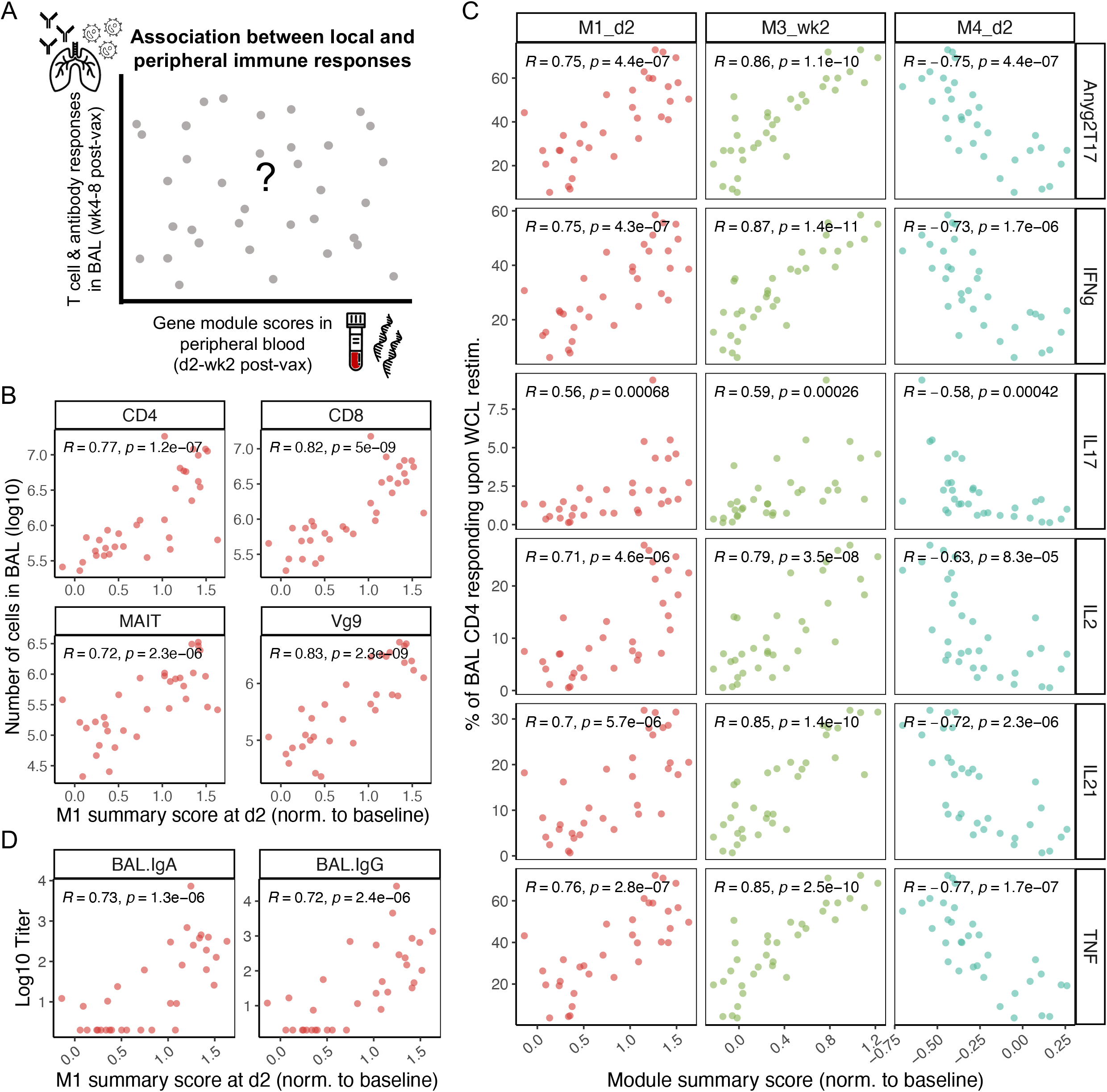
Correlations between module scores in blood and adaptive immune responses in bronchoalveolar lavage (BAL). A) Correlations between module 1 scores at day 2 in blood and T cell counts (log10) at week 4 in BAL. B) Correlations between module 1 scores at day 2 post-vaccination, module 3 scores at week 2 post-vaccination, or module 4 scores at day 2 post-vaccination and the frequency of antigen-specific CD4 T cells in BAL at week 8 post-vaccination. Antigen-specific CD4 T cells are defined as the frequency of BAL CD4 T cells expressing a given cytokine or any combination of cytokines (Anyg2T17) upon *ex vivo* restimulation with *Mtb* antigens. C) Correlations between module 1 scores at day 2 in blood and IgA or IgG antibody titers (log10) at week 4 in BAL or plasma. Pearson correlation coefficients and corresponding p-values are shown. All module scores were normalized to each animal’s baseline.

Indeed, innate module 1 scores in blood on day 2 post-vaccination were highly correlated with CD4, CD8, MAIT, and Vγ9 T cell numbers, identified by flow cytometry in BAL four weeks post-vaccination (Pearson correlation coefficient *r* ≥ 0.72, p≤2.3e-06; **Fig 2B**). Module 1 scores on day 2 were also correlated with numbers of B cells, NK cells, and DCs (*r* ≥ 0.61, p≤5.3e-04) in BAL, but not macrophages (**Fig S2A)**. Importantly, module 1 scores on day 2 were associated with the frequency of antigen-responsive CD4 T cells in BAL (*r* ≥ 0.56, p≤6.8e-04), identified by flow cytometry as CD4 T cells expressing IFNγ, IL-2, IL-17, IL-21, and/or TNF in response to *Mtb* restimulation eight weeks post vaccination (**Fig 2C**). The frequency of antigen-responsive CD4 T cells in BAL at 8 weeks post-vaccination was also strongly foreshadowed in blood by module 3 scores at week 2 (*r* ≥ 0.59, p≤2.6e-04) and module 4 score on day 2 (|*r*| ≥ 0.58, p≤4.2e-04; **Fig 2C)**. However, none of the modules were significantly associated with the frequency of antigen-responsive CD8 T cells in BAL **(Fig S2B, Fig S3)**. Finally, module 1 and module 4 scores at day 2 post-vaccination were strongly correlated with IgA and IgG titers in BAL at four weeks post vaccination (|*r*| ≥ 0.72, p≤2.4e-06) **(Fig 2D).**

We next examined whether any of the seven IV BCG-induced modules were associated with protection following *Mtb* challenge, despite having been discovered without considering the outcomes following *Mtb* challenge. Eighteen of 34 (53%) macaques were protected against TB (12 high-dose and 6 low-dose), as determined by fewer than 100 total colony forming units (CFUs) of *Mtb* upon necropsy. A higher dose of IV-BCG was associated with increased likelihood of protection (univariate odds ratio for a 10-fold increase in dose=3.602, p=0.013), but sterile protection (0 *Mtb* CFU) was nonetheless observed across all dose groups (**Fig 3A)**. We first assessed potential correlates of protection in the discovery subset of high-dose IV BCG recipients. When we modeled module activity over time post-vaccination using generalized estimating equations (**Methods)**, none of the seven modules differed significantly over time by protection outcome, likely due to small sample size. However, module 1 scores at day 2 post-vaccination were elevated in high-dose recipients that were protected following *Mtb* challenge compared to those that were not protected (Wilcoxon one-sided p=0.052; **Fig 3B**). The difference in module 1 scores at day 2 by protection outcome was conserved and more pronounced in the validation subset of low-dose recipients (Wilcoxon one-sided p=1.1e-4; **Fig 3B)**. Of note, although module 1 induction varied by dose, module 1 scores at day 2 remained significantly associated with protection outcome after adjusting for dose (p<0.001; **Table S2; Methods)**. Although other modules were associated with protection in low-dose IV BCG recipients (e.g., module 2 at day 2, module 3 at week 2, and module 4 at day 2), they were not associated with outcome in high-dose recipients (**Fig S4)**, suggesting that they may be important but insufficient for protection.

**Figure 3.**
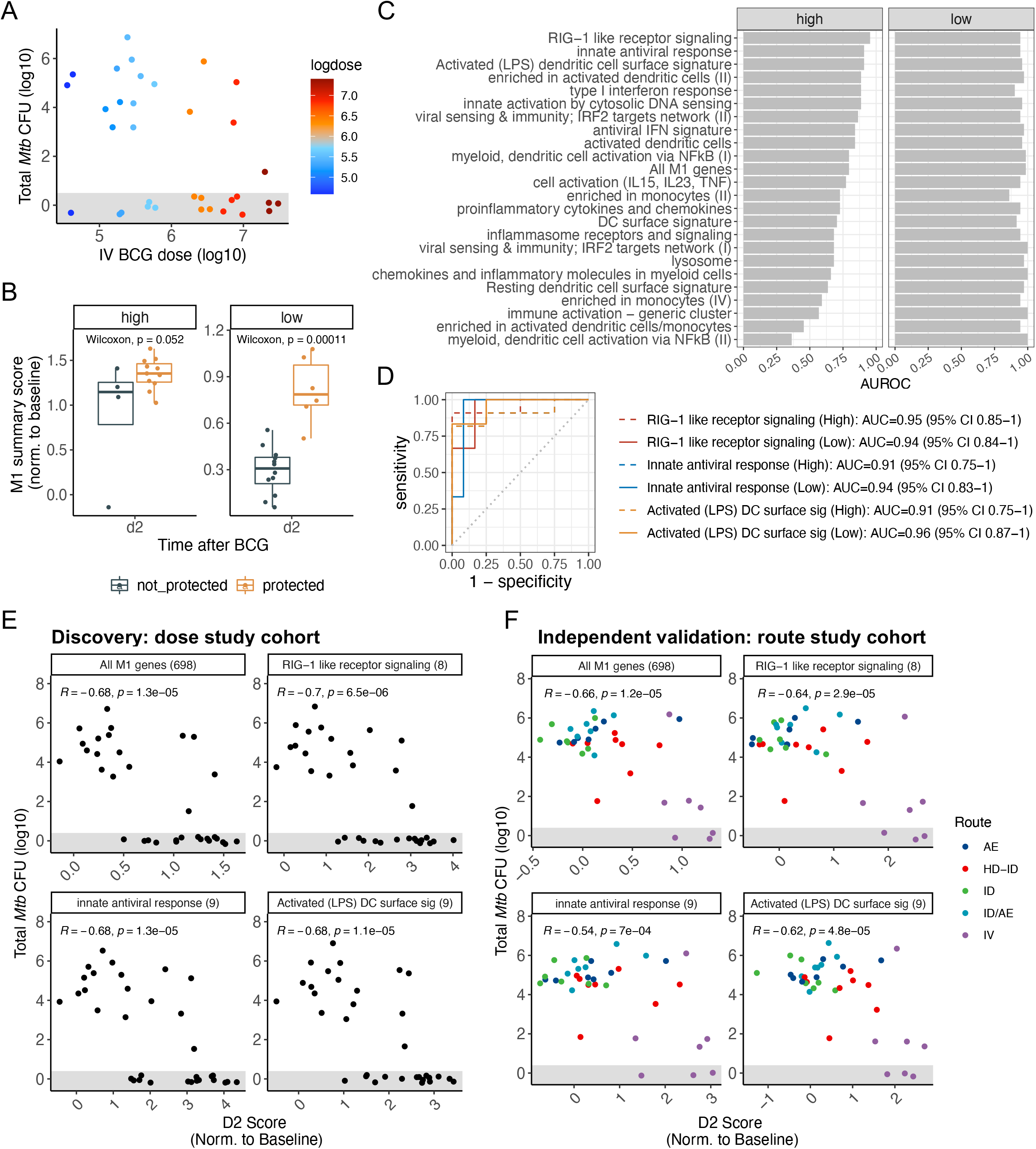
Innate module 1 is a reliable correlate of BCG-induced protection. A) Total *Mtb* colony-forming units (CFU) upon necropsy by IV BCG dose received. Animals with total *Mtb* CFU under 10^2^ (beneath the horizontal dashed line) are considered protected. B) Module 1 scores at day 2 for high-dose or low-dose IV BCG recipients, stratified by protection outcome following *Mtb* challenge. C) All immune pathways enriched in module 1 and their accuracy at day 2 in predicting protection following challenge among high- or low-dose recipients. AUROC, area under the receiver operating characteristic (ROC). D) ROC curves for the most accurate module 1 subpathways in high-dose recipients (dashed lines) and low-dose recipients (solid lines). E-F) Correlations between *Mtb* burden (log10 CFU) upon necropsy and day 2 scores of module 1 or its sub-pathways in the E) dose study cohort or F) route study cohort (AE, aerosol; HD-ID, high-dose intradermal; ID, intradermal; ID/AE, intradermal and aerosol; IV, high-dose intravenous.). Pearson correlation coefficients and corresponding p-values are shown. The number of genes in each set is shown in parentheses to the right of the set name. All scores were normalized to each animal’s baseline. Data points are jittered to reduce overplotting; all points in the shaded grey area represent animals with no detectable CFUs.

Given the consistent, reproducible association of module 1 scores at day 2 with protection following *Mtb* challenge, we further analyzed this module. Module 1 was comprised of 698 genes and enriched for 24 different innate immune pathways (FDR <0.01) (**Table S1**). We hypothesized that different pathways represented in module 1 would vary in their association with protection. To test this, we computed signature scores using the day 2 expression of module 1 genes in each enriched pathway, and we assessed the accuracy of these scores in predicting protection following challenge using AUROCs. Indeed, compared to all genes in module 1 which had an AUROC of 0.795 (95% CI, 0.520-1) in high-dose recipients, sub-pathways in module 1 had AUROCs ranging from 0.363 to 0.955 (95% CI: 0.852-1) in high-dose recipients, with the RIG-I-like receptor (RLR) signaling pathway exhibiting the highest accuracy (**Fig 3C-D**). Of note, all sub-pathways had high accuracy in low-dose recipients, with all but one pathway achieving AUROCs over 0.9 (**Fig 3C-D**). Strikingly, in addition to predicting binary protection outcome, day 2 scores of module 1 and its parsimonious sub-pathways were strongly correlated (|*r*| ≥ 0.59, p≤2.8e-04) with post-challenge *Mtb* burden and with the number of granulomas upon necropsy (**Fig 3E, Fig S5A)**. Importantly, these findings were validated in the independent cohort of macaques receiving BCG via various routes of administration (**Fig 3F, Fig S5B)**.

## DISCUSSION

Correlates of vaccine-mediated protection against TB will inform the development of more efficacious vaccines. Lung-localized immune responses can be useful for identifying correlates and understanding mechanisms of protection, but BAL samples are more difficult to obtain. Therefore, there is a critical need to identify blood-based correlates that can be more easily measured and scaled. However, TB vaccine studies to-date have failed to identify correlates in blood that have sufficient predictive accuracy. In this study we leveraged a non-human primate model of IV BCG vaccination followed by *Mtb* challenge to identify correlates of protection in the blood transcriptome. In one cohort where macaques were randomized to receive varying doses of IV BCG, increasing dose was associated with both stronger transcriptional responses to vaccination and higher levels of protection following *Mtb* challenge. We identified a gene module (module 1), predominantly comprised of innate immune pathways, whose score on day 2 post-vaccination was strongly associated with protection following challenge, even after adjusting for dose. Individual pathways within module 1, consisting of 8-9 genes each, robustly predicted protection outcomes following challenge with AUROCs >= 0.9 in both high- and low-dose IV BCG recipients. Notably, we validated module 1 and its sub-pathways as robust correlates of protection in an independent cohort of macaques receiving BCG through IV and other routes. Furthermore, module 1 scores on day 2 post-vaccination were highly correlated with functional T, B, and NK cell responses in the BAL at 4-8 weeks post-vaccination, and inversely correlated with the number of granulomas and *Mtb* CFUs in lung following challenge. Collectively, these findings shed light on the previously underappreciated early innate response to vaccination in peripheral blood, which validated across dose groups and in an independent cohort as an accurate, accessible predictor of protection against TB in macaques.

Within module 1, RIG-I-like receptor (RLR) signaling emerged as the pathway with the highest accuracy in predicting protection following challenge in high-dose IV BCG recipients. Strikingly, this accuracy was validated in low-dose IV BCG recipients and in an independent cohort of macaques vaccinated through different routes. RLRs are part of a cytosolic nucleic acid sensing system that activates host defense pathways against viral and intracellular bacterial infections, including *Mtb* [25]. However, unlike *Mtb*, BCG may fail to adequately activate RLRs due to its sequestration in immature phagosomes, precluding efficient lysosomal degradation and antigen presentation [26; 27]. Interestingly, a recent study showed that a dinucleotide agonist of RIG-I increased antigen presentation and IFNβ secretion by BCG-infected macrophages and DCs *in vitro* and enhanced BCG-mediated protection against *Mtb* challenge in a mouse model [28]. Accordingly, in the present study, several DC activation and interferon signaling pathways were also induced by IV BCG and associated with protection following challenge across dose groups. Together, these findings suggest that IV BCG-mediated protection may stem in part from increased activation of RLRs, induction of type 1 IFN signaling, and antigen presentation. Future research should investigate the mechanisms by which IV BCG achieves this early response to illuminate targets for novel adjuvants to improve BCG efficacy. Moreover, given that type 1 IFN signaling has been implicated in other settings as a correlate of risk for TB infection and disease [29; 30], future studies should elucidate the timing- and context-dependent roles of type 1 IFNs in mediating protection or susceptibility.

We identified additional elements of the transcriptional response that are likely important for—albeit not sufficiently predictive of—BCG-mediated protection, as evidenced by their association with protection in low-dose but not high-dose IV BCG recipients. For instance, module 2, which was also enriched for innate immune cell (monocyte, DC, and neutrophil) pathways, was more transiently upregulated than module 1 but may contribute to priming a subsequent local response. Furthermore, the early downregulation of T and NK cell pathways in module 4 on day 2 post-vaccination may in part reflect lymphocyte homing to secondary lymphoid organs for activation. The ensuing induction of CD4 T cell proliferation pathways in module 3 at weeks 2-4 following vaccination likely also reflects a critical step in establishing a robust lung-resident memory CD4 T cell response.

Our findings are critical to consider alongside those from a separate analysis of flow cytometry, Luminex, and complete blood count (CBC) data from the same dose cohort (Darrah *et al*, submitted). In that analysis, adaptive responses in BAL were better predictors of protection following challenge than responses in blood. However, in the present study, we found that several modules (M1, M3, and M4) measured at day 2 or week 2 postvaccination were highly correlated with T cell, NK cell, and antibody responses in BAL at week 4-8 post-vaccination. In addition, our early innate transcriptional signatures in blood predicted protection following *Mtb* challenge with comparable accuracy to the best-performing adaptive BAL features from the separate analysis by Darrah et al. (submitted). Therefore, in peripheral blood, transcriptional signatures may provide better correlates of protection than flow cytometry, Luminex, or CBC data, and may be adequate surrogates for lung-resident memory responses when BAL samples are unavailable.

Interest in the innate immune response to BCG vaccination has grown in recent years but has been largely directed toward BCG-induced ‘trained immunity’, or *de facto* innate memory [31]. These studies have primarily profiled later post-vaccination timepoints to demonstrate the longevity of BCG-induced changes in innate cells [32; 33; 34; 35; 36]. One study in rhesus macaques of a cytomegalovirus-based TB vaccine identified an innate blood transcriptional signature measured approximately 56 weeks post-vaccination, immediately before *Mtb* challenge, that was associated with post-challenge outcomes [13]. In contrast, to our knowledge, this study is the first to establish that blood transcriptional signatures induced *early* (within two days) post-vaccination are robustly correlated with protection against TB. This phenomenon has been shown for vaccines against other infections, including Ebola virus [18], influenza [16], and yellow fever [23]. Furthermore, to date, the field has focused on adaptive responses for identifying correlates of protection against TB [7; 37; 38; 39], which are usually profiled weeks or months after vaccination. Therefore, the early response to TB vaccination and its relevance for vaccine-mediated protection remains understudied.

Of the few studies that have examined the blood transcriptome at this early timepoint following *in vivo* vaccination with BCG or a TB vaccine candidate [40; 41; 42; 43], none have included subsequent TB outcomes. However, one study found a rapid increase of circulating neutrophils in mice and humans within 24 hours of BCG vaccination; in mice the increase in neutrophil count was required and sufficient for protection against polymicrobial sepsis, indicating that the early response to BCG may also predict non-specific BCG-mediated protection [41]. The neutrophil signal diminished by day 4 post-vaccination, further illustrating the time-restricted nature of certain responses to vaccination that may be overlooked in studies that only include later timepoints. Two other studies found early blood transcriptional responses induced by vaccine candidates M72/AS01E (in humans) [40] or H56/CAF01 (in mice) [42] that were similar to those we identified; namely, at 1-2 days post-vaccination they found upregulation of many of the same innate pathways as those enriched in module 1 of the present study, as well as downregulation of NK and T cell pathways. These overlapping findings in other species vaccinated with distinct subunit vaccines and adjuvants, delivered through non-IV routes (intramuscular or subcutaneous), suggest that certain elements of the blood transcriptional response may be conserved across diverse TB vaccine types and routes. It remains to be determined whether the association of these responses with protection is also generalizable.

A strength of our study is the systems-based approach that enabled comprehensive profiling of all aspects of the peripheral response to IV BCG vaccination, rather than only prespecified targets of interest. Moreover, our analyses included direct comparisons between peripheral transcriptional responses (blood) and local functional responses (BAL), adding to the limited understanding of the relationship between blood and lung biomarkers [44]. Our methodology was also intentionally designed for the challenges of high-dimensional data and a small sample size. Namely, we did not have sufficient statistical power to conduct gene-by-gene multivariate analyses with time, dose, protection, and their interactions as explanatory variables. Instead, we used a “discovery and validation” framework and divided the cohort by dose to minimize the confounding effects of dose while still ensuring our results were conserved across dose groups. After identifying significant DEGs in high-dose recipients, we conducted all subsequent analyses using gene modules in order to reduce dimensionality and avoid being prone to false discoveries or overfitting. Finally, we further validated the robustness of our findings in an independent cohort of macaques.

This study has several limitations. First, dose was a strong confounder, affecting both responses to vaccination and protection outcomes. Our methods sought to circumvent this as described above, but there were nonetheless striking differences between low- and high-dose recipients in the protection-associated response. Future studies should identify a single, intermediate dose that confers 50% protection for a more controlled correlates analysis. Next, IV BCG-mediated protection in rhesus macaques may differ from protection conferred by other vaccines, through other routes, and for other species. However, the overlap between our findings and those from other non-IV BCG studies in mice and humans suggests that some aspects of the protective response may be generalizable and may at least inform strategies to improve BCG efficacy for humans. Finally, our results were limited by the lower resolution of bulk RNA-sequencing. A priority for future research is to perform single cell-based analyses in order to elucidate cell type-specific responses and identify more granular correlates.

Altogether, our results provide new insights into the early innate response to TB vaccination, which we found to be a robust predictor of protection following *Mtb* challenge. While our findings in low-dose IV BCG recipients implicate other components that may be important for protection, such as T and NK pathways at later timepoints, these were not sufficient to explain differing outcomes following challenge in high-dose recipients. Instead, only the strength of select innate responses two days post-vaccination, such as the activation of RLRs and DC pathways, could accurately predict protection following challenge across dose groups and in an independent cohort of macaques. These findings underscore the critical need for additional research on the early responses required to prime long-lasting protective immunity against TB. This research could enable improved TB vaccine and adjuvant design and facilitate efficient evaluation of existing and future vaccine candidates.

## ACKNOWLEDGMENTS

We thank Hong Zheng, Michele Donato, Madeleine Scott, and Jason Andrews for valuable advice and discussions. This work was supported by the Intramural Research Program of the Vaccine Research Center at the National Institute of Allergy and Infectious Diseases (NIAID); the Bill and Melinda Gates Foundation (OPP1113682 to PK, Aeras to JLF, and BMGF INV-020435 to JLF); NIAID grants 1U19AI109662 (PK), U19AI057229 (PK), and 75N93019C00071 (JLF); Department of Defense contracts W81XWH-18-10253 and W81XWH1910235 (PK); the Ralph & Marian Falk Medical Research Trust (PK); the Knight-Hennessy Scholars Program (YEL); and the National Science Foundation Graduate Research Fellowship Program (YEL). The funders had no role in study design, data collection and analysis, decision to publish, or preparation of the manuscript.

## AUTHOR CONTRIBUTIONS

Conceptualization, Y.E.L., P.A.D., M.R., J.L.F., R.A.S., and P.K.; Methodology and Formal Analysis, Y.E.L.; Investigation, Y.E.L, P.A.D., J.J.Z., M.K., F.L., D.C.D., and P.M.; Writing – Original Draft, Y.E.L., Writing – Review & Editing, Y.E.L., P.A.D., J.J.Z., M.K., F.L., D.C.D., P.M., M.R., J.L.F., R.A.S., and P.K.; Supervision, M.R., J.L.F, R.A.S., and P.K.

## METHODS

### Study design and analytical approach

To identify correlates of BCG-induced protection, we utilized the “dose” cohort, in which thirty-four rhesus macaques of Indian origin were vaccinated intravenously (IV) with varying doses of BCG as detailed in a separate manuscript (Darrah et al, submitted). Macaques received between 3.9×10^4^ and 2.5×10^7^ CFUs of IV-BCG (Danish strain 1331), which we dichotomized into a binary variable of high or low dose (greater than or less than 10^6^ CFUs) for our main analyses, although we also performed additional analyses of dose as a continuous variable. Whole blood was collected for bulk RNA sequencing at baseline (four weeks before vaccination) and two days, two weeks, four weeks, and twelve weeks following vaccination. Twenty-four weeks following vaccination, macaques were challenged by bronchoscope with a low dose of barcoded *Mtb* Erdman (mean 12 CFU) and euthanized 12 weeks later, or at the humane end point, for analysis of disease burden (Darrah et al, submitted).

Due to the limited sample size and high-dimensional nature of the data, we were underpowered to adjust for dose and its interactions with time and protection outcome in multivariate models. Therefore, we took a training-and-validation approach, performing discovery analyses in high-dose recipients and validating our findings in low-dose recipients to ensure generalizability by dose. Specifically, we first identified vaccine-induced genes and modules in high-dose recipients without considering outcomes following challenge. We then assessed which modules were associated with protection following challenge and validated putative correlates in low-dose recipients. We also examined associations between blood transcriptional responses and local immune responses in the lung (BAL) measured using multi-parameter flow cytometry and Luminex assays as outlined in a separate manuscript (Darrah et al, submitted).

We further validated our findings in an independent cohort of macaques from the “route” cohort for which immune and outcome data are published [12]. Briefly, thirty-six macaques received BCG vaccination through different routes: aerosol (AE, n=7), intradermal (ID, n=7), high-dose intradermal (HD-ID, n=8), combined aerosol and intradermal (AE/ID, n=7), and intravenous (IV, n=7). IV-vaccinated macaques in the route cohort received 5×10^7^ CFUs of BCG (high-dose). Macaques were challenged with *Mtb* six to ten months following BCG vaccination and euthanized 11-15 weeks following challenge or at humane endpoint. For this cohort, we analyzed bulk RNA sequencing data from whole blood samples collected at baseline and two days, two weeks, and twelve weeks following vaccination.

### RNA processing and sequencing

Whole blood was collected in PAXgene blood RNA tubes (Qiagen) and stored at −80C for batch processing at end of study. RNA was extracted using the PAXgene Blood RNA Tube kit (PreAnalytiX) as instructed. Globin mRNA was removed using GLOBINclear Kit (Life Technologies) and remaining mRNA concentration and quality was measured on an Agilent Bioanalyzer using an Agilent nano 6000 kit. Illumina-ready libraries were generated using NEBNext Ultra II RNA Preparation reagents (New England BioLabs). The barcoded Illumina-ready libraries were sequenced using paired-end 151-base protocol on either a NovaSeq 6000 (Illumina) or a HiSeq 4000 (Illumina).

### Transcriptomic analysis

#### Data processing

We trimmed reads with Cutadapt and performed quality control checks with FastQC before aligning to the Macaca mulatta genome (mmul_10) using kallisto. We removed one sample from the route cohort with fewer than 2000 total sequences. We removed genes in the globin family, genes that were unannotated, and genes with fewer than 5 reads in over two-thirds of all samples. We performed hierarchical clustering to check for outliers and removed one sample that clustered separately from all other samples. This resulted in 167 samples from the dose discovery cohort and 144 samples from the route validation cohort for analysis. We applied DESeq’s variance stabilizing transformation to generate normalized expression values for visualization. All data processing was performed independently for the dose study and route study cohorts.

| Time after BCG | Baseline | 2 days | 2 weeks | 4 weeks | 12 weeks | Total |
| --- | --- | --- | --- | --- | --- | --- |
| Number of samples (dose study cohort) | 33 | 33 | 34 | 33 | 34 | 167 |
| Number of samples (route study cohort) | 36 | 36 | 36 | NA | 36 | 144 |

#### Identification of BCG-induced genes and modules in high dose IV BCG recipients

We identified genes that were differentially expressed at each timepoint following vaccination, relative to baseline, in high dose IV-BCG recipients from the “dose” discovery cohort. Using DESeq2, we employed a multi-factor design with animal ID to account for between-subject variability at baseline. We utilized approximate posterior estimation for log2 fold change (LFC) shrinkage [45] and applied significance thresholds of |LFC| ≥ 1.5 and adjusted p-value ≤ 0.1. To identify coordinated gene modules induced by BCG vaccination, we performed weighted gene correlation network analysis (WGCNA) [46] with the union of genes that were differentially expressed at one or more timepoints following vaccination. We chose the soft thresholding power of 9, which achieved a scale-free topology R^2^ of 0.85. We set a minimum module size of 50 genes and merged similar modules whose distance was less than 0.2. One hundred and ninety-one genes remained unassigned. We summarized the expression pattern of each of the final seven modules using a summary score computed as the geometric mean expression of all genes in each module, normalized to each animal’s baseline. For one macaque that was missing a baseline sample, we imputed baseline gene expression values using the median baseline expression across all other animals.

#### Pathway analysis

To determine the immune pathways enriched in each module, we performed over-representation analysis using the hypergeometric test, with blood transcription modules (BTMs) as the reference gene set database [23]. Human gene names in the BTM database were converted to rhesus macaque Ensembl gene IDs using *biomaRt* [47]. We applied a significance threshold of FDR ≤ 0.01 and excluded “to be annotated” (TBA) BTMs. For sub-module BTM “scores”, we computed the geometric mean expression of module genes in each significantly enriched BTM, normalized to baseline for each animal.

#### Correlation between peripheral and local response

Immune responses in the bronchoalveolar lavage (BAL) after IV BCG vaccination were measured as described separately (Darrah et al., submitted). In the present study, we focused on B- and T-cell responses in BAL that were previously shown in the route cohort to be higher after IV BCG vaccination, compared to AE or ID BCG at the same dose. We compared module expression in blood with the following immune markers in BAL: IgA and IgG antibody titers against *Mtb* whole cell lysate; CD4, CD8, Vγ9 gamma delta, and MAIT T cell counts; and antigen-specific memory CD4 and CD8 T cell responses (production of IFNγ, IL-2, IL-17, IL-21, TNF upon restimulation with *Mtb* purified protein derivative (PPD)). Modeling module expression over time by protection outcome. To test whether module expression over time following vaccination was significantly different by protection outcome, we utilized generalized estimating equations (GEE), which is an extension of generalized linear models that accounts for within-subject (animal) correlation for repeated measures data. We fit separate models for each module and for high-dose or low-dose recipients. Baseline, day 2, and week 2 timepoints were included. Module summary score was the response variable. Explanatory variables were time (in days), time^2^, time^3^, binary protection outcome, and interactions between each time term and protection outcome. We included quadratic and cubic terms based on empirically observed trends in module activity over time and on a goodness-of-fit metric (quasilikelihood under the independence model criterion or QIC). We computed 95% confidence intervals and p-values using robust standard errors. We considered module expression over time as significantly different by protection outcome if any of the interaction terms were significant with p≤0.05.

#### Non-linear model to assess correlates of protection adjusted by dose

To determine whether module expression was still correlated with protection after adjusting for dose, we fit a non-linear least squares model using a generalized logistic function akin to a dose-response curve with the following formula:

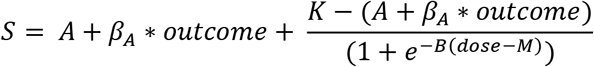

Where *S* is the module score at a given time (normalized to baseline), *A* is the lower asymptote which varies by *outcome* (binary protection outcome) by a fitted coefficient of *β_A_, K* is the upper asymptote, *B* is the growth rate, and *M* is the dose midpoint or curve inflection point. We report *β_A_* and its corresponding p-value as an indicator of the dose-adjusted estimate for how module expression differs between animals that were or were not protected following challenge.

#### ROC curves

To compare the accuracy of peripheral transcriptional responses versus local immune responses to vaccination in predicting protection following challenge, we computed area under the receiver operating characteristic (ROC) curves (AUROCs). For blood transcriptional responses, we used the geometric mean scores defined above for modules and BTMs within a module. We computed separate AUROCs for high dose and low dose recipients in order to assess generalizability by dose.

## SUPPLEMENTARY FIGURES

**Figure S1. Module activity in macaques from the route study cohort.** Summary scores of all seven modules over time following vaccination in the route study cohort, where macaques were BCG-vaccinated via different routes. AE, aerosol; HD-ID, high-dose intradermal; ID, intradermal; ID/AE, intradermal and aerosol; IV, high-dose intravenous. Shown are module scores for individual macaques (thin grey lines) and the median per group (thick colored lines).

**Figure S2. Correlations between module scores in blood and immune responses in BAL.** A) Correlations between module 1 scores at day 2 post-vaccination and number (log10) of B cells, NK cells, iNKT cells, macrophages, mDCs, or pDCs in BAL. All BAL cell counts were measured four weeks post-vaccination except NK cell counts which were measured eight weeks post-vaccination. pDC cell count measurements were missing for nine (26%) animals. B) Correlations between module 1 scores at day 2 post-vaccination, module 3 scores at week 2 post-vaccination, or module 4 scores at day 2 post-vaccination and the frequency of antigen-specific CD8 T cells in BAL at week 8 post-vaccination. Antigen-specific CD8 T cells are defined as the frequency of BAL CD8 T cells expressing a given cytokine or any combination of cytokines (Anyg2T17) upon *ex vivo* restimulation with *Mtb* whole cell lysate. Pearson correlation coefficients and corresponding p-values are shown. All module scores were normalized to each animal’s baseline.

**Figure S3. Complete correlation matrix for module scores in blood and immune responses in BAL or plasma.** Pairwise pearson correlation coefficients (*r*) are shown between summary scores for each module at each timepoint and various immune responses of interest. Correlation p-values were adjusted using the Benjamini-Hochberg method; significant correlations at a false discovery rate threshold of 0.01 are indicated with an asterisk. BAL, bronchoalveolar lavage; Ag-spec, antigen-specific; Anyg2T17, expressing any combination of IFNγ, IL-2, TNF, IL-17. Antigen-specific CD4 and CD8 T cell responses are defined as the frequency of BAL CD4 or CD8 T cells expressing a given cytokine upon *ex vivo* restimulation with *Mtb* whole cell lysate eight weeks post-vaccination. BAL counts of various cell types and antibody titers are on a log10 scale and were measured four weeks post-vaccination, with the exception of NK cell counts which were measured eight weeks postvaccination. pDC cell count measurements were missing for nine (26%) animals. All module scores were normalized to each animal’s baseline.

**Figure S4. Module activity stratified by IV BCG dose and outcome following challenge.** Summary scores (normalized to baseline) of all seven modules over time following vaccination, stratified by IV BCG dose and whether macaques were protected or not protected following *Mtb* challenge. The dotted horizontal line at y=0 indicates no change from baseline.

**Figure S5. Correlations between the granuloma burden following challenge and module 1 pathways at day 2 post-vaccination.** Correlations between the number of granulomas (log10) upon necropsy and day 2 scores of module 1 or its sub-pathways in the A) dose study cohort or B) route study cohort. Pearson correlation coefficients and corresponding p-values are shown. The number of genes in each set is shown in parentheses to the right of the set name. All scores were normalized to each animal’s baseline. Data points are jittered to reduce overplotting; all points in the shaded grey area represent animals with no detectable CFUs.

**Figure S6. ROC curves for T cell responses in BAL post-vaccination as predictors of protection following challenge.** A) ROC curves for absolute counts of CD4, CD8, MAIT, or Vg9 T cells in BAL at week 4 post-vaccination. B) ROC curves for antigen-specific CD4 T cell responses in BAL at week 8 post-vaccination. Solid and dashed lines represent low- and high-dose IV BCG recipients, respectively. AUC, area under the ROC curve; CI, confidence interval; Anyg2T17, expressing any combination of IFNγ, IL-2, TNF, or IL-17.

## SUPPLEMENTARY TABLES

**Table S1. Enriched blood transcriptional pathways in each module.** Only gene sets with an adjusted p-value (padj) <= 0.01 are shown. Original size indicates the number of genes in each set. Filtered size indicates the number of genes in each set that were measured in this study. Relevant genes indicates the number of genes in each set that were present in each module.

**Table S2. Fitted parameter values from non-linear model adjusted for dose.**

