## Supplementary figures and images for "Blood Transcriptional Correlates of BCG-Induced Protection Against Tuberculosis in Rhesus Macaques"

### Figure S1

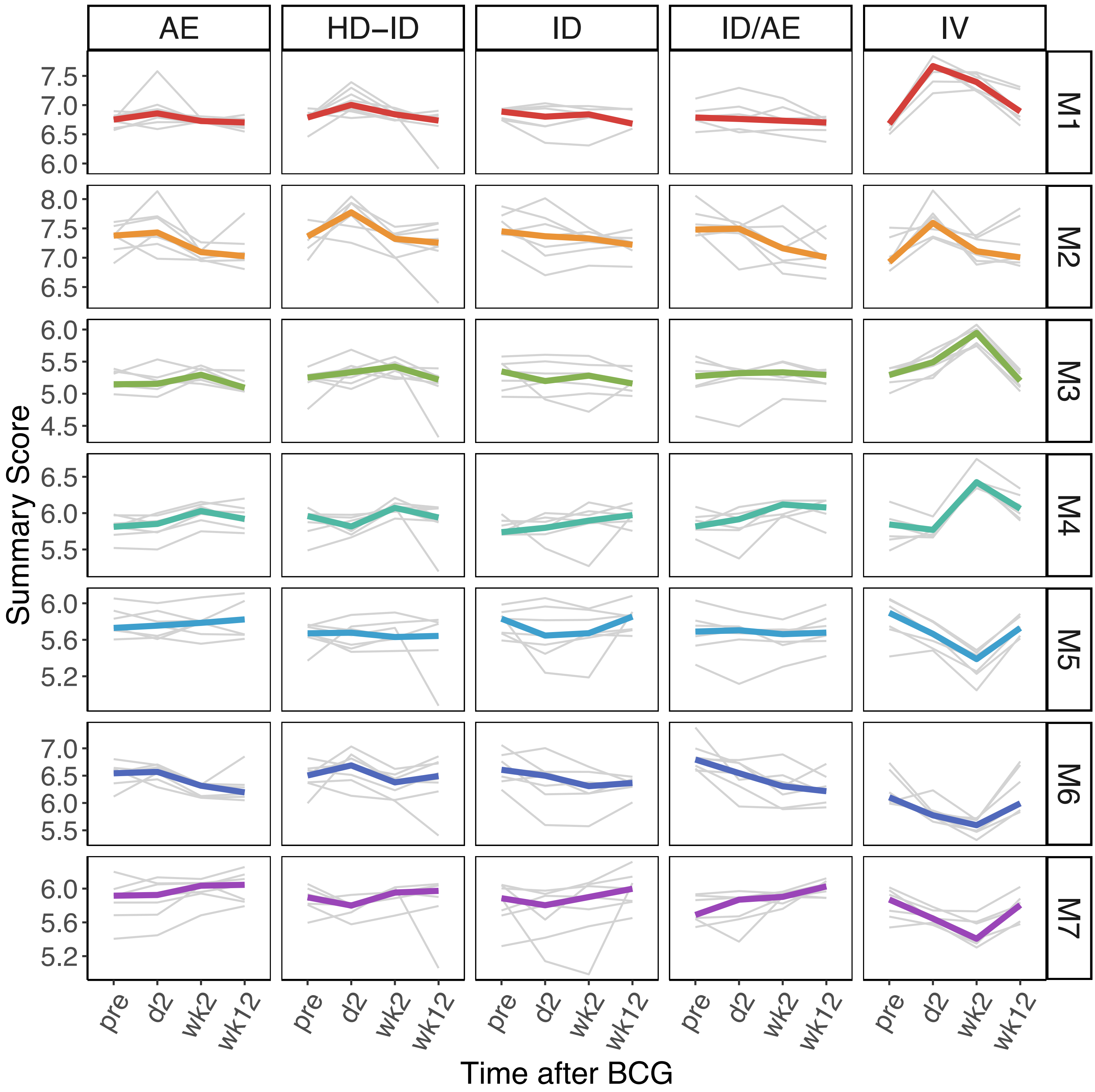

### Figure S2

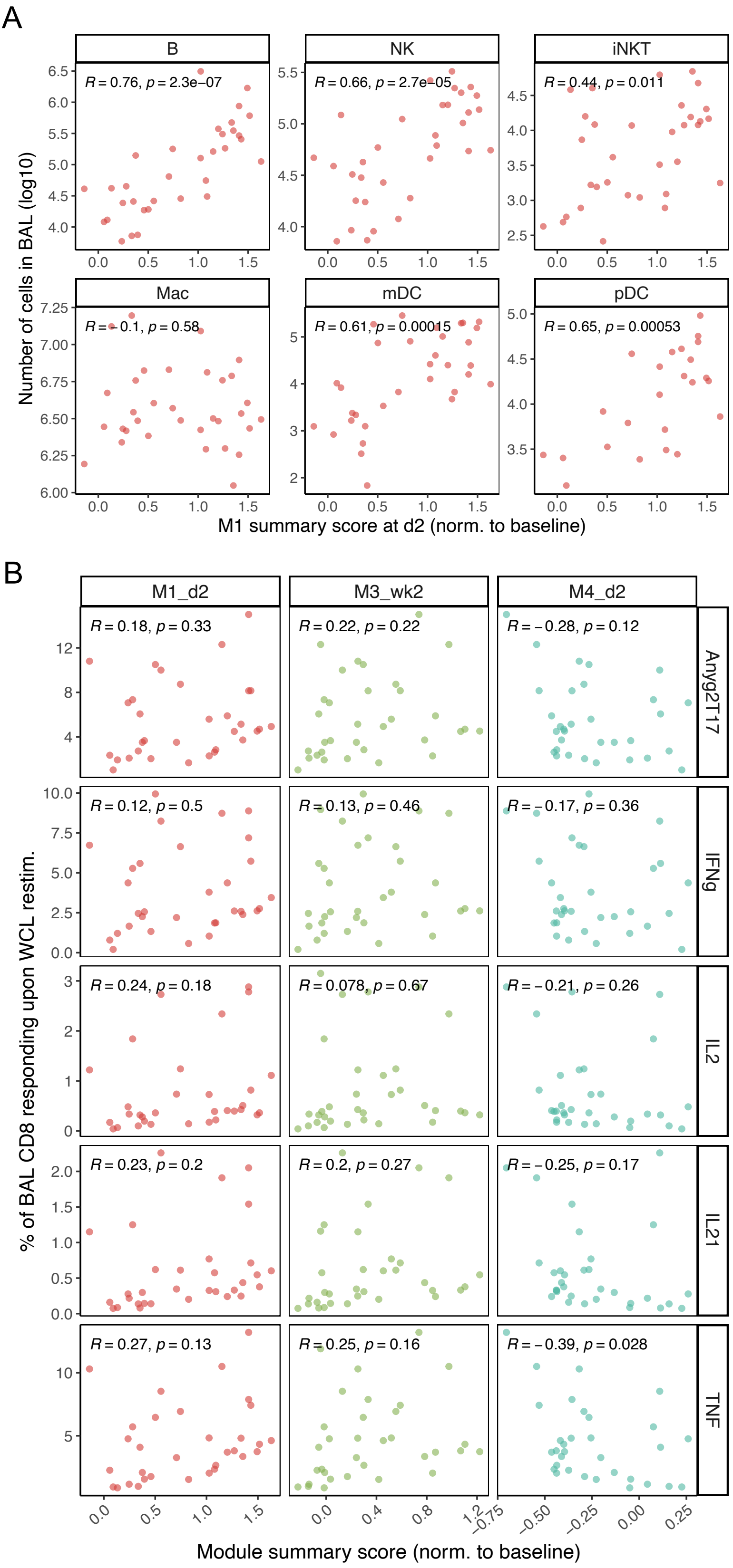

### Figure S3

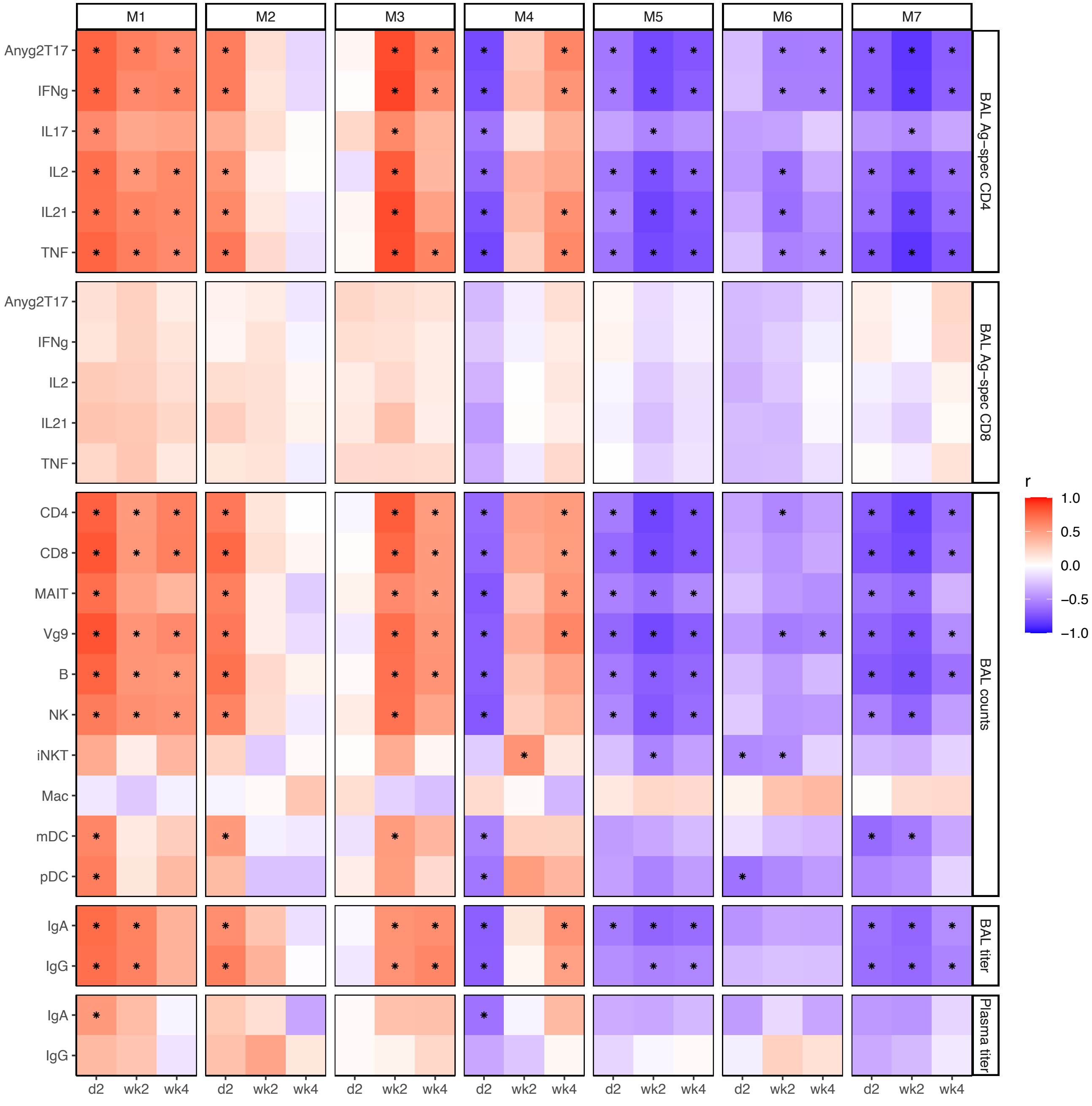

### Figure S4

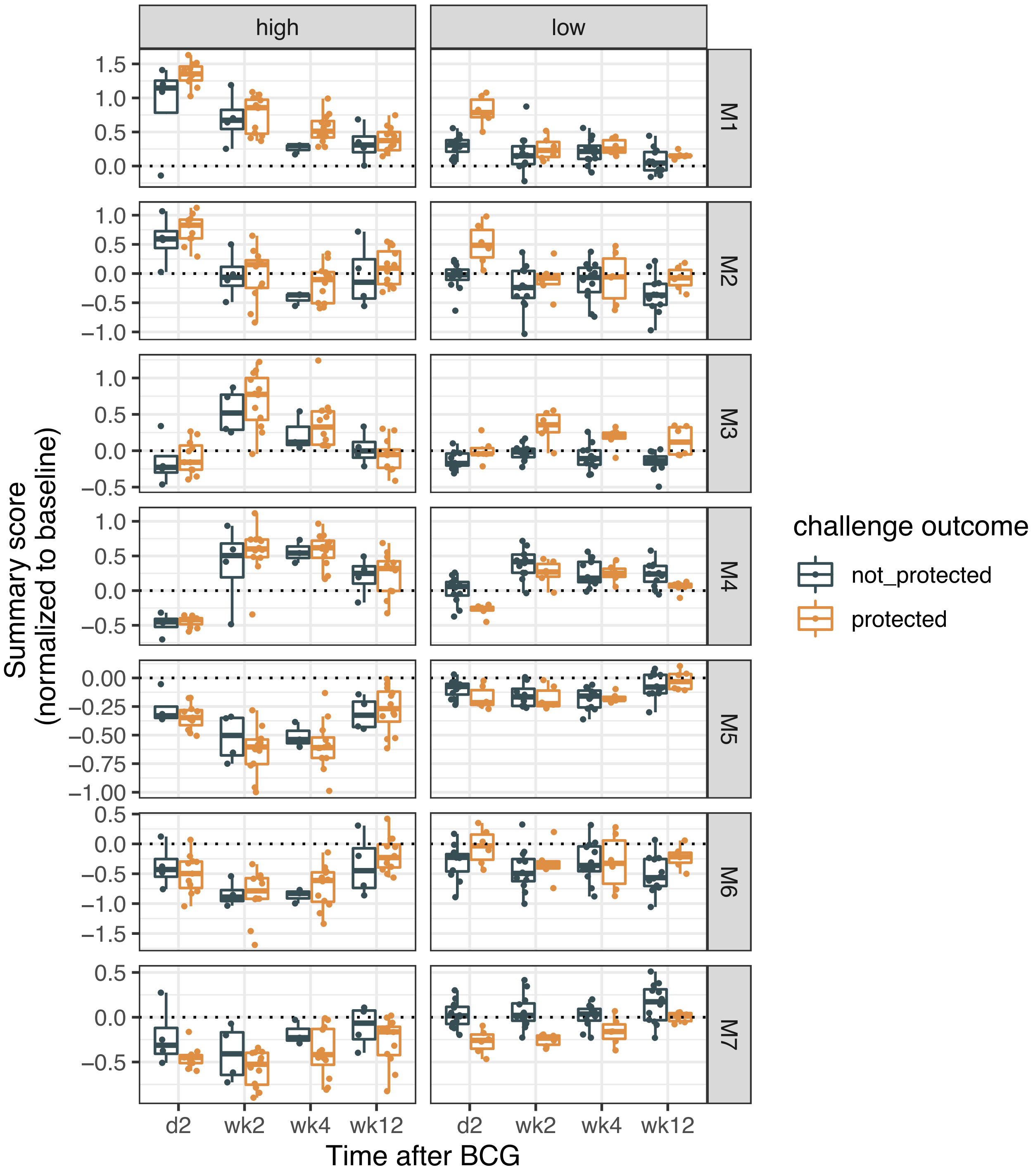

### Figure S5

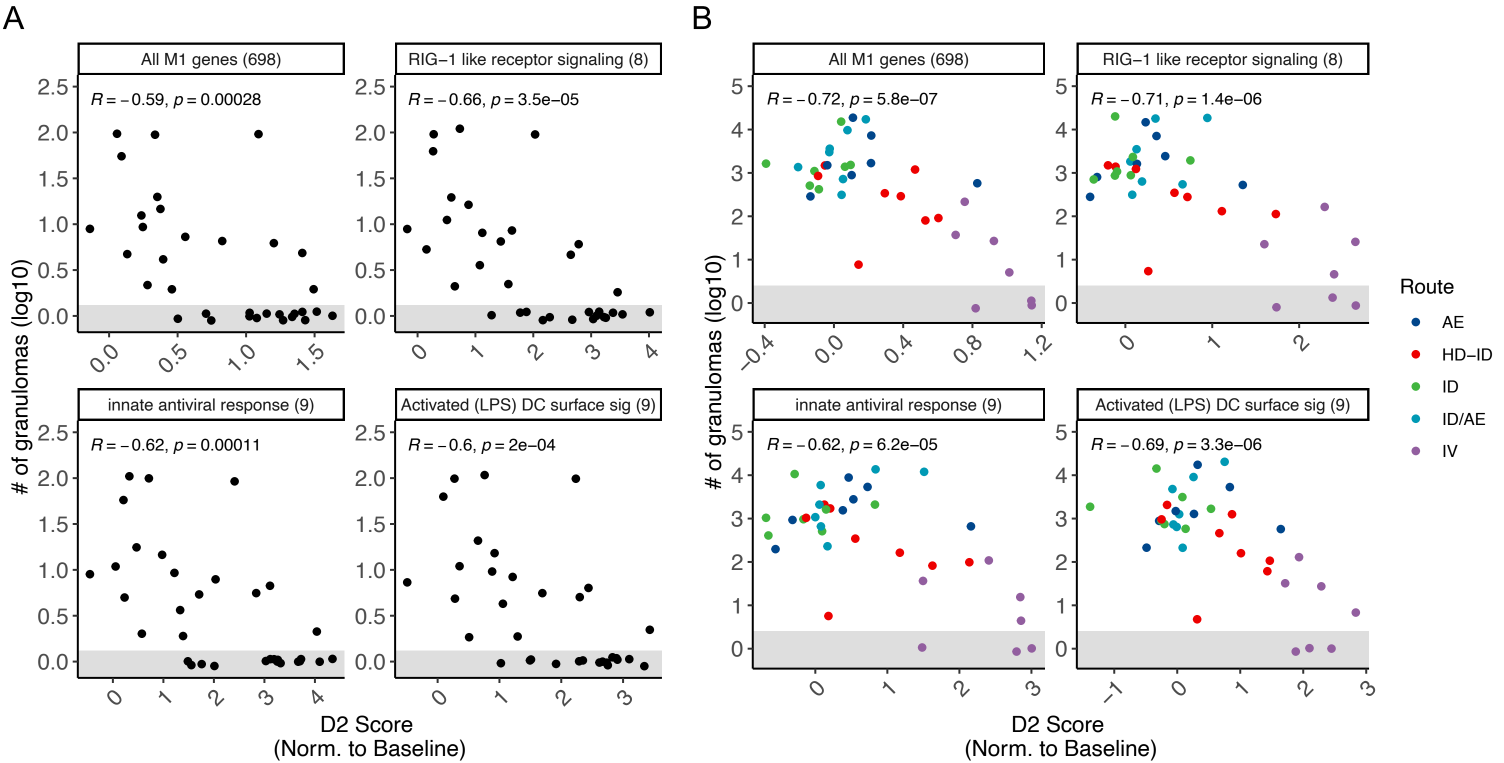
